# The Microglial *Trem2* R47H Alzheimer’s Disease Risk Variant Does Not Detectably Alter Hippocampal Synaptic Transmission in Young Mice

**DOI:** 10.1101/2025.08.22.671714

**Authors:** Abdulaziz Aljawder, Hedaythul Choudhury, Damian M. Cummings, Jack Wood, Frances A. Edwards

## Abstract

The rare R47H variant of the microglial *TREM2* gene increases Alzheimer’s disease (AD) risk, but its effects on early hippocampal circuitry remain unclear. *Trem2* has been reported to be important for microglial-dependent refinement of excitatory synaptic circuits in development. We hypothesized that loss of TREM2 function might result in changes in circuitry due to reduced microglial ability to prune inactive synapses and that this could predispose individuals to later disease. We compared basal synaptic transmission and synaptic density in 3-week-old WT and homozygous *Trem2* R47H knockin mice, using *in vitro* patch clamp recordings and Bassoon-Homer1 immunohistochemistry. Basal synaptic activity, miniature excitatory events, and evoked release probability were unchanged. Synapse density in CA1 and CA3 showed substantial between-mouse variability but no evidence of genotype or regional differences. This variable CA1 synapse density was positively correlated with CA1 miniature excitatory event frequency. Although between-mouse variability in synaptic density could mask subtle changes, these data indicate that early effects of *Trem2* R47H in these mice are not sufficient to produce a consistent change in early postnatal hippocampal excitatory synapse density or basal synaptic transmission under these conditions. This supports the view that *TREM2*-linked AD risk may act primarily through context-dependent mechanisms that become relevant with ageing, neuroinflammatory challenge, or AD pathology.

## Introduction

Triggering receptor expressed on myeloid cells 2 (TREM2) is a microglial cell surface receptor ^1,2^. Homozygosity for the rare R47H partial loss-of-function variant in the *TREM2* gene causes Nasu-Hakola disease, which is characterized by early-onset dementia, multifocal bone cysts, and may also lead to epilepsy^3–5^. Additionally, a range of experimental approaches have identified that heterozygosity for the R47H missense variant in *TREM2* is one of the strongest genetic risk factors for sporadic AD, conferring up to a fourfold increase in disease risk^6,7^. Most studies of the effects of *Trem2* R47H in AD models have concentrated on its role in altering the interaction of microglia with plaques^8–10^.

However, early-life events can also influence later susceptibility to neurodegenerative disorders (reviewed in^11,12^). Studies of such susceptibility to early-life influence are often related to changes in immune function with most reports relating to environmental factors such as perinatal hypoxia^13^. Microglia play an essential role in pruning inactive synapses, a process thought to be critical for the maturation and refinement of functional neuronal circuits^14–16^. This role in refinement of circuits has been suggested to be TREM2-dependent and, if this is the case, then it is possible that *TREM2* mutations, such as R47H, alter the synaptic network during development, and that this predisposes carriers to AD. Although several studies address this question, most have relied on knockout of *Trem2* in cultures or, in some cases, young animals and the results have been conflicting, variously suggesting both increases and decreases in synaptic density and function^16–18^. As TREM2 influences the activity of a pathway of proteins involved in phagocytosis and lysosomal function^19^, at least in older animals, knockout of *Trem2* may result in complex interactions and homeostatic responses that may not relate to the normal function of its receptor, and this may explain the complex outcomes seen under slightly different conditions.

In a study in 5-6-month-old mice heterozygous for the *Trem2* R47H mutation, Das et al. (2023) reported changes in synaptic networks in both wild type (WT) mice and in a rapidly progressing model of early AD (*App^NL-G-F/NL-G-F^*knockin mice)^20^. Specifically, the *Trem2* R47H expression increased susceptibility to kainate-induced seizure activity, which the authors attributed to elevated synapse density observed in the cortex of both WT and *App* knockin mice carrying the *Trem2* mutation. No significant difference was observed in synaptic density in the hippocampus at this age^20^.

Despite these studies, research on the electrophysiological impact of the *Trem2* R47H variant remains limited. In 6-8 weeks old homozygous R47H rats, Schaffer-collateral-CA1 evoked responses were larger, had reduced long-term potentiation (LTP), and the same authors subsequently found a TNF-α-dependent weakened inhibitory signaling onto CA1 cells^21,22^. Studies on a different *Trem2* loss-of-function mutation, *Y38C*, in 6-month-old mice, have also demonstrated a significant impairment in LTP^23^.

As microglia have an important role in refining synaptic circuits in early development, understanding how genetic variants, such as *TREM2* R47H, affect microglia-mediated synaptic development may provide crucial insights into additional mechanisms by which this variant increases AD risk.

In this study of 3-week-old mice, preweaning, we use whole-cell patch clamp recordings to assess hippocampal synaptic function and use immunohistochemistry to investigate density of synapses in both the CA1 and CA3 subfields of the hippocampus. We found little evidence of change in either synaptic activity or density in the mice carrying the *Trem2* R47H mutation. However, we observe a surprising variability in synaptic density between mice, which is not dependent on genotype.

## Results

To assess the effects of the *Trem2* R47H mutation on synaptic function in young mice, whole-cell patch clamp recordings were performed from CA1 pyramidal neurons in acute hippocampal slices prepared from mice homozygous for the *Trem2* R47H variant and from WT controls preweaning, at 18–22 days of age. It is important to note that translational differences between human and murine *TREM2* R47H variants in these mice result in a decrease in expression of TREM2 of around 70%^8^; an effect not observed in humans^19,24^. Thus, the experiments in this study represent not only a putative loss of function due to the R47H mutation but also partial knockdown of *TREM2* expression. However, the general principle of early effects of microglial dysfunction would apply in both cases and indeed more widely across other forms of neurodegeneration.

### Comparison of synaptic currents in CA1 region with and without the *Trem2* R47H mutation

#### Overall frequency of spontaneous synaptic currents is not altered by the Trem2 R47H variant

Spontaneous postsynaptic currents (sPSCs) recorded in artificial cerebrospinal fluid (aCSF) reflect the overall synaptic activity impinging on the recorded pyramidal cells. This consists of a mixture of spontaneous excitatory postsynaptic currents (sEPSCs) and spontaneous inhibitory postsynaptic currents (sIPSCs).

Analysis of sPSCs in CA1 pyramidal neurons from *Trem2* R47H and WT mice revealed no significant differences between genotypes in event frequency, median amplitude, or decay time-constant (τ), suggesting that the *Trem2* R47H variant does not impact overall synaptic transmission (Figure 1). However, total spontaneous activity could mask differential changes in excitatory and inhibitory currents.

**Figure 1.**
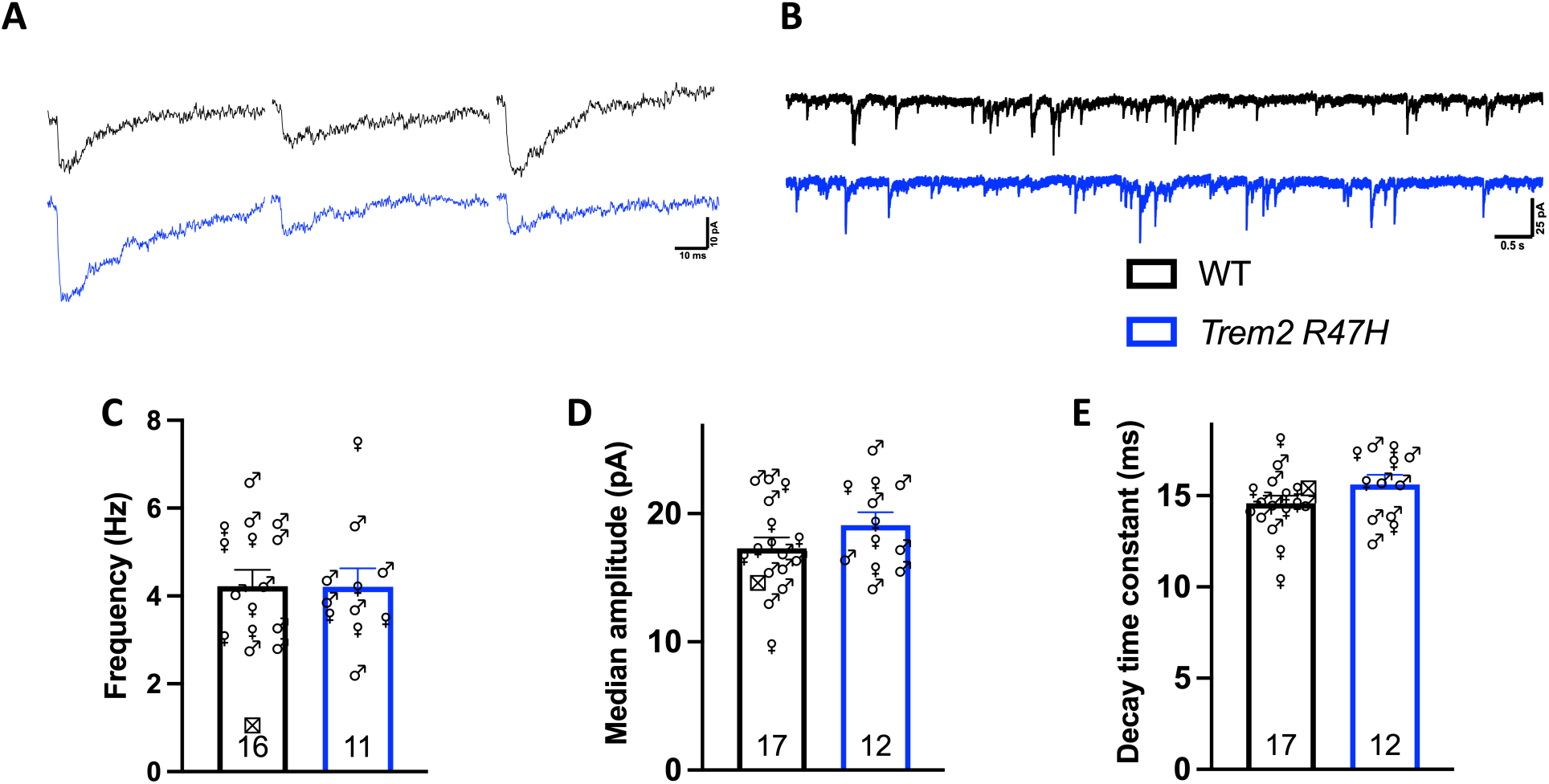
The *Trem2* R47H variant does not affect combined spontaneous postsynaptic currents. (A) Representative sPSC current traces recorded from CA1 pyramidal neurons in WT and *Trem2* R47H mice. Scale bars are indicated. (B) Extended 10 s traces of sPSCs in WT and *Trem2* R47H neurons. (C–E) Quantification of sPSC frequency (C), median amplitude (D), and τ (E) across genotypes. Data are presented as mean + SEM. Statistical comparisons were performed using unpaired *t*-tests. Numbers displayed on the bars indicate sample size, and symbol shapes denote the sex of the mouse from which each recording was made. Crossed square symbols indicate mice for which sex information was unavailable.

#### Separating sEPSCs from sPSCs suggests no change in either excitatory or inhibitory currents

To isolate sEPSCs, GABA^A^ receptor-mediated inhibition was pharmacologically applied using gabazine (6 μM, Figure 2A). Analysis of sEPSCs recorded from CA1 pyramidal neurons revealed that event frequency and median amplitude were not normally distributed and so nonparametric Mann–Whitney U tests were applied. An unpaired Student’s *t*-test was used to compare τ between the cohorts. No significant differences in frequency, median amplitude or τ were observed between the genotypes (Figure 2B–E).

**Figure 2.**
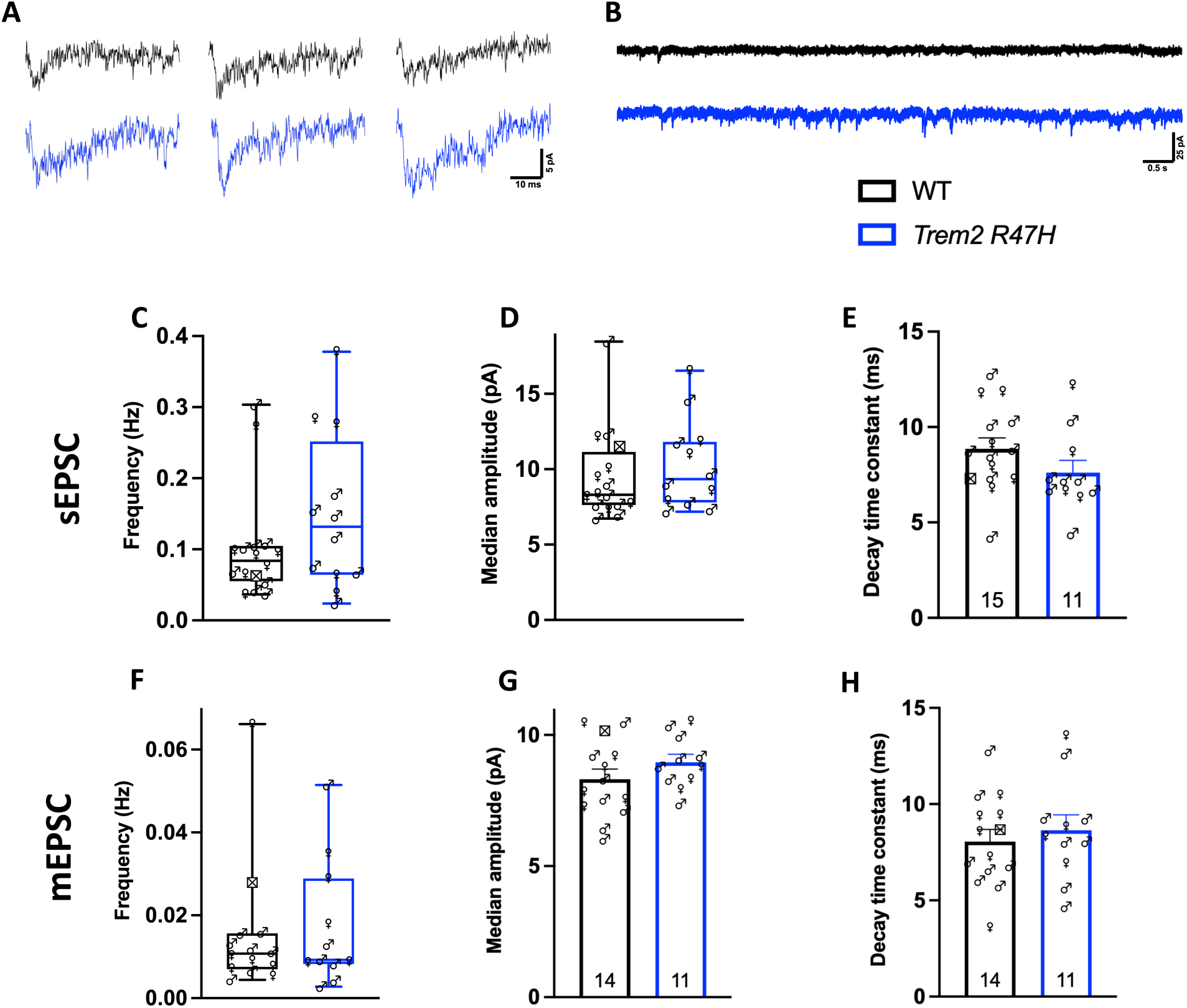
The *Trem2* R47H variant does not change sEPSC or mEPSC synaptic measures. (A) Representative sEPSC traces from CA1 pyramidal neurons in WT and *Trem2* R47H mice. (B) 10 s continuous recordings of sEPSCs from WT and *Trem2 R47H* mice. (C–E) Quantification of sEPSC frequency (C; *n* = 15 WT, *n* = 11 *Trem2* R47H), median amplitude (D; *n* = 15 WT, *n* = 11 *Trem2* R47H), and τ (E). (F–H) Quantification of mEPSC frequency (F, *n* = 14 WT, *n* = 11 *Trem2* R47H), median amplitude (G), and τ (H). Data are presented as median and IQR in C, D and F, and as mean + SEM in E, G and H. Statistical comparisons were performed using Mann–Whitney U tests in C, D and F, and unpaired *t*-tests in E, G and H. Numbers displayed on the bars of bar graphs indicate sample size, and symbol shapes denote the sex of the mouse from which each recording was made. Crossed square symbols indicate mice for which sex information was unavailable. Scale bars are indicated as appropriate.

The fact that, in the presence of gabazine, the frequency of synaptic currents decreased from approximately 4 Hz to 0.1 Hz on average suggests that the combined spontaneous activity is very largely (> 95%) dominated by inhibitory synaptic currents. This is only an approximation because there would be some interaction between the two populations. Nevertheless, as there was no significant change in the overall sPSC, and no significant difference in the frequency of the small proportion of sEPSCs contributing to this, we can conclude that there are unlikely to be changes in sIPSC frequency or the other variables measured.

#### Isolating miniature excitatory postsynaptic events from sEPSCs shows no change in Trem2 R47H

To isolate miniature excitatory postsynaptic currents (mEPSCs), 1 μM tetrodotoxin was added to the extracellular solution containing gabazine, thereby blocking action potential-dependent synaptic activity. Analysis of mEPSCs recorded from CA1 pyramidal neurons revealed no significant differences between *Trem2* R47H and WT mice in event frequency (Mann–Whitney U test), amplitude (*t*-test), or τ (*t*-test; Figure 2).

Together, these findings suggest that the *Trem2* R47H mutation does not detectably change sEPSCs, sIPSCs and mEPSCs in CA1 pyramidal cells in the postnatal hippocampus.

#### Trem2 R47H mutation does not alter the probability of glutamate release in young mice

As the frequency of spontaneous activity is dependent on the number of synapses as well as their activity in terms of release probability and axonal excitability, it is possible that changes in synaptic pruning could be masked by compensatory changes in release probability. We thus went on to assess the strength of synaptic connections and release probability by evoking EPSCs. In order to compare release probability between the two genotypes, we recorded responses to paired-pulse stimulation at inter-stimulus intervals of 50 and 100 ms. The paired-pulse ratio (ratio of the amplitude of the second response to the first response at short stimulus intervals; PPR) is inversely related to neurotransmitter release probability (Figure 3)^25^.

**Figure 3.**
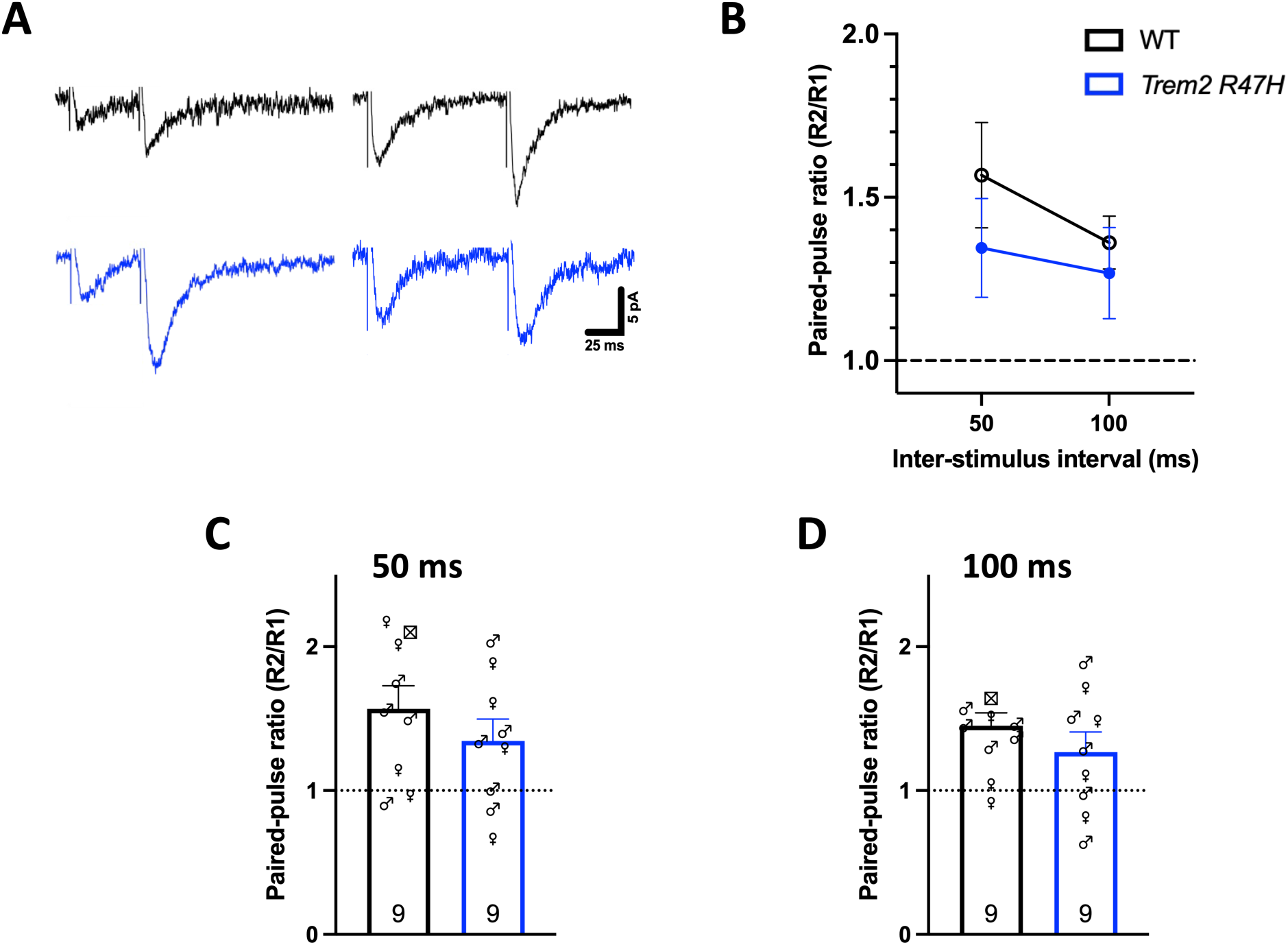
The *Trem2* R47H variant does not alter presynaptic glutamate release probability. (A) Representative traces of evoked excitatory postsynaptic currents recorded from CA1 pyramidal neurons in WT and *Trem2* R47H mice at each inter-stimulus interval. Stimulus artifacts are attenuated for clarity. Scale bars are indicated. (B) Summary of paired-pulse ratios across the 2 inter-stimulus intervals (50, and 100 ms; mean ± SEM). Statistical analysis was performed using 2-way repeated-measures ANOVA. (C,D) Quantification of paired-pulse ratios at each inter-stimulus interval: 50 ms (C), and 100 ms (D). Data are presented as mean + SEM. Statistical comparisons were performed using unpaired *t*-tests. Numbers displayed on the bars indicate sample size, and symbol shapes denote the sex of the mouse from which each recording was made. Crossed square symbols indicate mice for which sex information was unavailable.

Two-way repeated-measure ANOVA showed no significant main effect of inter-stimulus interval or *Trem2* genotype, and no significant interaction (Figure 3B). Quantification of PPRs at each inter-stimulus interval revealed no significant differences between *Trem2* R47H and WT mice according to unpaired *t*-tests, suggesting that the variant does not impact presynaptic release dynamics in the developing hippocampus (Figure 3 C,D).

Of note, when males and females were analysed separately, no significant WT versus *Trem2* R47H differences were detected for sIPSC, sEPSC, or mEPSC frequency, median amplitude, or decay kinetics (data not shown).

Overall, the electrophysiological results suggest that at early ages, shortly prior to weaning, basal synaptic transmission and release dynamics are not altered by the R47H mutation of *Trem2* and/or decreased TREM2 levels.

### Comparison of synaptic density in CA1 and CA3 regions with and without the *Trem2* R47H mutation

If the *Trem2* R47H variant impairs appropriate synaptic pruning during development, it would be surprising that this has little, if any, effect on basal synaptic transmission in young mice. However, both in the case of spontaneous activity and when evoking action potentials using extracellular stimulation of axons in the above sections, we would tend to select for axons that had relatively high release probability. Axons with low release probability could therefore be missed. Considering synaptic pruning targets relatively inactive synapses, that is those with a low release probability, it is possible that retention of such synapses would be difficult to detect with patch clamp techniques. Inappropriate retention of synapses that were largely inactive in early development could still result in changes later in life as synaptic plasticity alters synaptic strength.

To investigate synaptic density alterations in the hippocampus, immunohistochemistry was performed on brain sections from the fixed hemisphere of WT and *Trem2* R47H mice.

Presynaptic terminals were labelled with anti-Bassoon antibodies, postsynaptic densities with anti-Homer1 antibodies, and cell nuclei with DAPI. Super-resolution Airyscan imaging followed by 3D reconstruction in Imaris software was used for synaptic quantification (Figure 4A–C).

**Figure 4.**
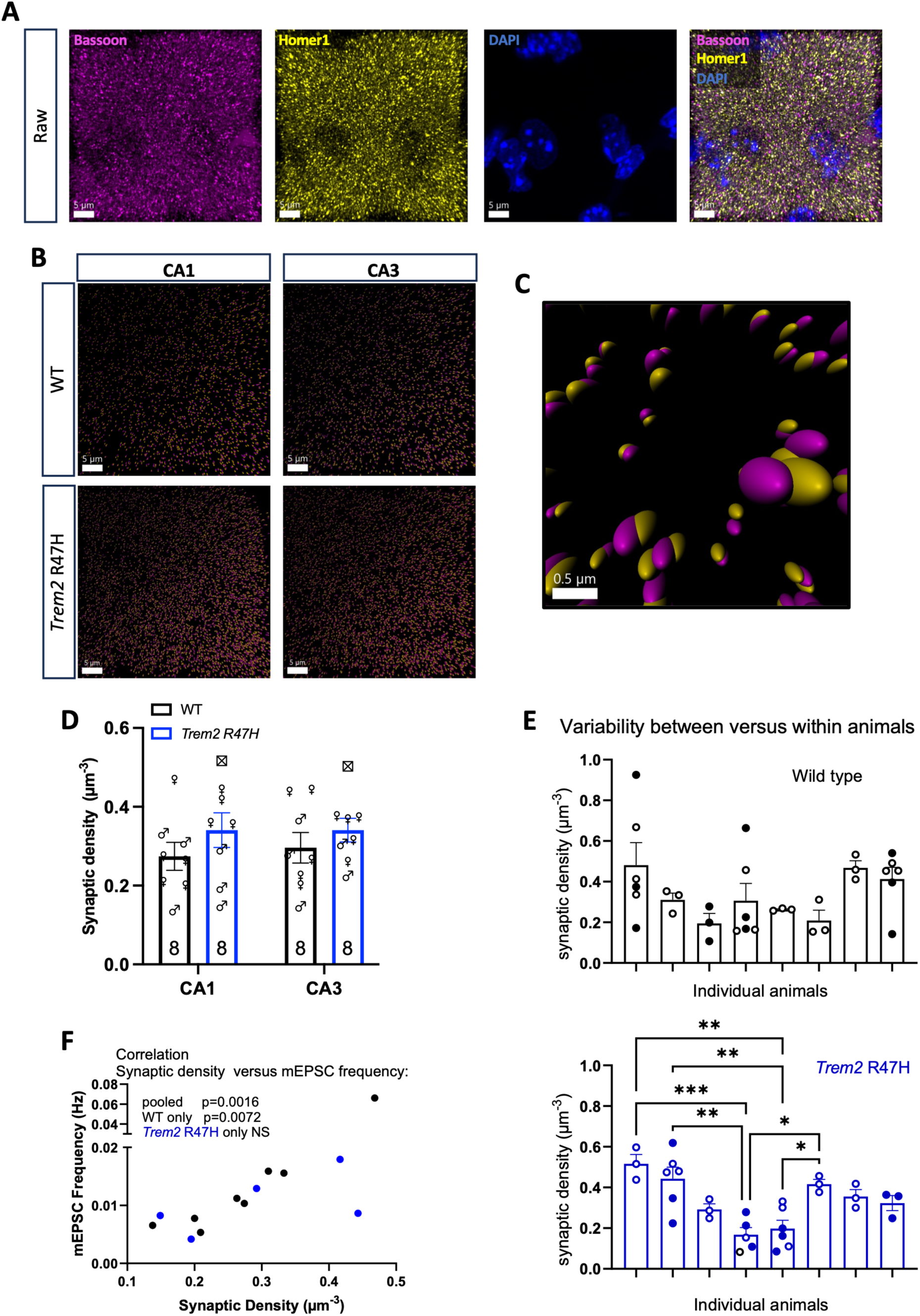
*Trem2* R47H does not alter excitatory synapse density in early postnatal hippocampal CA1 and CA3 regions. (A) Representative confocal Airyscan-processed images of regions of interest in the hippocampus showing immunofluorescent labelling of presynaptic Bassoon (magenta), postsynaptic Homer1 (yellow), and nuclear DAPI (blue). (B) Representative Imaris three-dimensional reconstructions of colocalized synaptic puncta in hippocampal CA1 and CA3 regions from WT and *Trem2* R47H mice. (C) A higher magnification of the colocalization in B. (D) Quantification of synaptic density (colocalized puncta per µm³) in CA1 and CA3 regions. Statistical analysis was performed using two-way repeated-measures ANOVA to assess the effects of genotype and hippocampal subregion. Data are presented as mean ± SEM. Numbers displayed on the bars indicate sample size, and symbol shapes denote the sex of the mouse from which each recording was made. Crossed square symbols indicate mice for which sex information was unavailable. (E) Comparison of variation between animals and variation between sections analyzed within each animal. Columns represent the mean synaptic density in individual animals. Symbols represent the synaptic density in individual sections within each animal. Closed and open symbols represent independent analyses in different sections by two different individuals blind to genotype. One-way ANOVA shows no significant difference between cells in WT mice but a highly significant difference between individual *Trem2* R47H mice (*P* < 0.0001). Tukey’s multiple comparisons are indicated in the figure for significant differences. (F) Correlation between CA1 excitatory synapse density (Bassoon-Homer1 colocalized puncta per µm³) and CA1 mEPSC frequency plotted for WT (black) and *Trem2* R47H (blue) mice. A positive correlation was seen between synaptic density and mEPSC frequency as shown on the graph for both data pooled across the genotypes (Spearman *r* = 0.80; *P* = 0.0016; *n* = 13) and for the data from WT animals alone (Spearman *r* = 0.88; *P* = 0.0072; *n* = 8). The data from the *Trem2* R47H mice showed no significant correlation (*n* = 5). Scale bars indicate x–y dimensions and correspond to 5 µm in panels A and B, and 0.5 µm in panel C. \**P* < 0.05; \*\**P* < 0.01; \*\*\**P* < 0.001.

Synaptic puncta were defined as Bassoon and Homer1 colocalized signals. Signals located within DAPI-labeled nuclear volumes were excluded from analysis to focus on extranuclear fluorescent signals. Two-way repeated-measure ANOVA revealed no significant effects of genotype or region (CA1 versus CA3), and no genotype-region interaction (Figure 4 B, D). These results indicate that the *Trem2* R47H variant did not alter synaptic density in either CA1 or CA3 regions of the hippocampus. Sex-stratified analyses of CA1 and CA3 synapse density showed no significant WT versus *Trem2* R47H differences in either males or females (data not shown).

#### Between-mouse variability in synaptic density and its relationship with frequency of mEPSCs

In the course of the above studies, it was notable that there was considerable variability between individual mice of the same genotype (Figure 4D). Although, in some cases, the synaptic density analyzed from different sections from an individual mouse showed considerable variation, generally the variability between individual animals was considerably greater than the variability between sections from the same animal, especially in the *Trem2* R47H group. One-way ANOVA revealed a highly significant difference in the synaptic density between individual animals in the *Trem2* R47H mice but no significant difference between WT mice (Figure 4E).

To determine whether the observed variation in anatomical synapse density was reflected in basal quantal excitatory transmission, CA1 Bassoon-Homer1 synapse density was compared to CA1 mEPSC frequency in all mice for which both measurements were available.

Associations were tested using Spearman’s rank correlation. CA1 synapse density showed a significant positive association with mEPSC frequency both when WT mice were tested alone (Spearman *r* = 0.8810, *P* = 0.0072, *n* = 8) or when both genotypes were pooled (Spearman *r* = 0.8802, *P* = 0.0016, *n* = 13; Figure 4F). There was no significant correlation when the results of mice carrying the *Trem2* R47H genotype were tested alone, but this may have been because insufficient mice were available for this comparison (*n* = 5).

The strong correlation between synaptic density and mEPSC frequency in the CA1 region suggests that the variability in synaptic density between mice is indeed a real phenomenon. Some of the variability could have arisen from positions of the sections along the hippocampal axis.

However, as 3-6 sections were taken from each animal for IHC, along the length of the hippocampus, with different randomly selected sections used for synaptic analysis, it is unlikely that the position of the section from dorsal to ventral had an influence. Although electrophysiological data was not available from the CA3 region, it seems that there is considerable variability between animals in the synaptic density in both the CA1 and CA3 regions.

## Discussion

The main finding of this study is that homozygous *Trem2* R47H did not produce a detectable change in the basal hippocampal synaptic measures examined at P18–P22. Spontaneous synaptic currents, isolated sEPSCs, mEPSCs, Schaffer-collateral PPRs, and Bassoon-Homer1 synapse density in CA1 and CA3 were not significantly different between *Trem2* R47H and WT mice. Because total spontaneous activity was dominated by inhibitory currents under the present recording conditions, the lack of change in overall sPSCs also argues against a large genotype effect on spontaneous inhibitory activity, although inhibitory synapses were not isolated. Therefore, any effect of *Trem2* R47H during early development is either too subtle to be detected by the present experiments, is restricted to synaptic features not measured here, or is offset by compensatory mechanisms associated with reduced TREM2 expression or function.

These findings help define the experimental conditions under which *Trem2* R47H produces detectable synaptic phenotypes. Previous studies reported altered Schaffer collateral-CA1 transmission, LTP, and inhibitory signaling in young homozygous *Trem2* R47H rats^21,22^, whereas the present study examined basal synaptic transmission in preweaning mice and did not assess LTP or inflammatory modulation. Similarly, increased synapse density and network hyperexcitability have been reported in older *Trem2* R47H mice, but those effects were most evident in cortex rather than hippocampus^20^. The absence of a detectable hippocampal phenotype here therefore suggests that *Trem2* R47H-associated synaptic changes may depend on age, species, brain region, disease context, or the physiological endpoint measured.

A notable feature of the dataset was the substantial between-animal variability in both synaptic activity and synaptic density. Although differences in synaptic density did not reach statistical significance for the WT mice, this probably reflected the greater within mouse variability in some animals which was less pronounced in the *Trem2* R47H mice. For synaptic activity, some of this variability could reflect differences in slice health or other experimental factors across recordings. However, where matched anatomical and electrophysiological data were available from the same mouse, CA1 Bassoon-Homer1 synapse density was strongly correlated with CA1 mEPSC frequency suggesting that structural variability was reflected in functional synaptic measures. This analysis suggested that the between-animal variation observed may reflect biologically meaningful differences in excitatory synaptic organization rather than only technical variability. The source of this between-mouse variability remains unclear, but it could reflect litter-level, maternal, or other early postnatal environmental factors. Since broad variability was present in both genotypes, the present data do not support a simple *Trem2* R47H-dependent shift in excitatory synapse density. However, the highly significant between-animal differences in synaptic density among the *Trem2* R47H mice could reflect different degrees of compensation for dysfunction in early synaptic pruning.

In conclusion, homozygous *Trem2* R47H did not detectably alter the measured indices of basal CA1 synaptic transmission, Schaffer collateral PPR, or CA1/CA3 excitatory synapse density by P18–P22. Future studies using longitudinal designs would be needed to determine whether the early between-animal variability observed in synaptic density predicts later behavioral, physiological, or disease-related outcomes.

## Limitations of the study

The present study has several limitations that warrant consideration. While the presynaptic marker Bassoon is a core component of the active zone at both excitatory and inhibitory synapses^26^, the postsynaptic marker Homer1 is predominantly localized to excitatory synapses, where it functions as a scaffolding protein within the postsynaptic density^27^. Consequently, our synaptic colocalization analysis did not investigate inhibitory synapses, limiting our ability to generalize the measured synaptic density across all synapse types.

For practical reasons the WT and *Trem2* R47H mice are not littermates as, with litter sizes of about 6, if we bred 2 heterozygous parents, only half of the litter would be either WT or *Trem2* R47H homozygous on average. We would thus frequently not get both genotypes occurring in the same litter. It was thus considered preferable to breed from WT or homozygous *Trem2* R47H parents to enable us to use all the mice. However, all other conditions were identical.

Furthermore, as outlined above, translational differences between human and murine *TREM2* R47H must be acknowledged. Specifically, Xiang et al. demonstrated (as further confirmed in the present lab) that in this mouse knockin model of the *Trem2* R47H, aberrant mRNA splicing leads to a decrease of about 60–70% in TREM2 expression, an effect not observed in human induced pluripotent stem cells-derived microglia or post-mortem brain^8,24^. Thus, the experiments in this study represent not only putative loss of function due to the R47H mutation but also partial knockdown of TREM2 expression. Thus, it remains possible that the decrease in TREM2 levels and the effect of the mutation work in opposite directions, screening changes that would occur with the R47H mutation alone. This seems unlikely as most evidence suggests that the R47H mutation results in decreased function of TREM2 and hence these differences would be expected to be complementary. It would nevertheless be of interest in future studies to utilize the recently established *Trem2* R47H mutant mouse line that does not undergo cryptic splicing^28^.

Additionally, the mice in this study are homozygous for *Trem2* R47H, which in principle may make them more relevant to Nasu Hakola disease than AD^3–5^. However, the general principle of early effects of microglial dysfunction would apply in both cases and indeed more widely across other forms of neurodegeneration.

## Resource Availability

### Lead Contact

Requests for additional information, resources, or reagents should be addressed to the lead contact, Frances A. Edwards.

### Materials Availability

The *Trem2* R47H mouse line generated in this study is subject to restrictions imposed by breeder material transfer agreements with The Jackson Laboratory (Bar Harbor, ME, USA).

### Data and Code Availability

All data from this study are available from the lead contact upon request. No original code is reported. Additional information necessary for data reanalysis can be obtained from the lead contact.

## Acknowledgements

This work was supported by grants from Alzheimer’s Research UK and the Cure Alzheimer’s Fund. We thank Haady Hajar for his assistance with mouse colony management in the Edwards Laboratory at University College London. A.A. is grateful to Arabian Gulf University for sponsoring his PhD studies at UCL; the data presented here form part of his doctoral research.

## Authors Contributions

Conceptualization: F.A.E. and A.A.

Methodology: A.A., H.C., D.M.C. and J.W.

Investigation: A.A., H.C.

Visualization: A.A., H.C., J.W. and D.M.C.

Supervision: F.A.E., D.M.C. and J.W.

Writing – original draft: A.A

Writing – review & editing: A.A., J.W., D.M.C., and F.A.E.

Funding: F.A.E.

## STAR★Methods

### Experimental model and study participants details

All procedures were performed in accordance with the UK Animals (Scientific Procedures) Act 1986. *Trem2* R47H knockin (Jackson Laboratory via MRC Harwell) and C57BL/6J WT mice were bred at University College London and used prior to weaning at P18–P22. All mice were housed within individually ventilated cages under a 12:12-hour light-dark cycle at the University College London Biological Services Unit. Food and water were provided *ad libitum*.

### Methods details

Pups were decapitated and brains bisected in ice-cold aCSF (in mM: 125 NaCl, 1.4 NaH₂PO₄, 26 NaHCO₃, 2.4 KCl, 20 glucose, 3 MgCl₂, 0.5 CaCl₂; pH 7.4, 310–315 mOsm). One hemisphere was fixed (4% PFA, 24 h, 4 °C) and cryoprotected (30% sucrose in phosphate-buffered saline (PBS)); the other was used for electrophysiology.

#### Electrophysiology

Acute transverse hippocampal slices were prepared from P18–P22 pups of both sexes according to Cummings et al^29^. Briefly, 300 μm-thick slices were prepared in ice-cold dissection aCSF, warmed to 35 °C, equilibrated in aCSF with increasing Ca²⁺ (0.5–2 mM) and decreasing Mg²⁺ (3–1 mM), and then maintained at room temperature for at least 40 minutes. Whole-cell voltage clamp recordings were performed at V_hold_ = -70 mV from CA1 pyramidal neurons identified using infrared differential interference contrast microscopy (Olympus BX50WI). Pipettes (4–6 MΩ) were filled with internal solution (in mM: 140 CsCl, 2 Mg-ATP, 10 EGTA, 5 HEPES; pH 7.4, ∼290 mOsm). Signals were amplified and low-pass filtered (2 kHz; Multiclamp 700B), digitized (10 kHz; Digidata 1322A) and then acquired within WinEDR (v3.2.7; University of Strathclyde). Series resistance (15–40 MΩ) was monitored throughout.

sEPSCs were recorded in aCSF with 6 μM gabazine. mEPSCs were recorded with 1 μM tetrodotoxin added. Events (≥3 pA for ≥2 ms, 10 ms deadtime) were detected using WinEDR and manually confirmed. Event frequency was calculated as 1/inter-event interval; amplitude was peak from baseline. Decay time constants (τ) were obtained by fitting a single exponential from peak to baseline.

PPRs were obtained by stimulating Schaffer collaterals in the presence of gabazine (100 μs constant-voltage pulses at 50 and 100 ms inter-stimulus intervals; ≥20 repeats were averaged for each inter-stimulus interval, with 10 s between successive stimulus pairs). Pipettes (aCSF-filled) were placed 100–350 μm from the recording site. Only monosynaptic responses (∼4 ms latency) were included in analyses; paired-pulse ratio was calculated as the amplitude of the second response/amplitude of the first response. All recordings and analyses were performed blind to genotype.

#### Immunohistochemistry

Standard protocols were followed to conduct immunohistochemical experiments^30^. Briefly, fixed frozen brain hemispheres embedded in 30% sucrose in PBS were sectioned using a sledge microtome (Leica SM2010R). Sections were cut perpendicular to the hippocampal longitudinal axis at a thickness of 30 μm, immediately transferred to PBS containing 0.02% sodium azide, and stored at 4 °C. Brain sections were permeabilized (0.3% Triton X-100 in PBS), blocked for non-specific binding (3% goat serum in 0.3% Triton X-100 in PBS) and then incubated with primary antibodies (1:500 guinea pig anti-Bassoon, Synaptic Systems, Catalog number 141318; and 1:250 chicken anti-Homer1, Synaptic Systems, Catalog number 160026) at 4 °C for 24 hours, and then with Alexa Fluor-conjugated secondary antibodies (1:500 goat anti-guinea pig AF 594, ThermoFisher, catalog number A-11076; and 1:500 goat anti-chicken AF 647, ThermoFisher, catalog number A-21449) for 2 hours at room temperature. Lastly, sections were mounted onto SuperFrost Plus™ adhesion slides using Fluoromount-G® mounting medium, and then stored at 4 °C.

#### Imaging

Two regions of interest were imaged per brain section: a defined area within the *stratum radiatum* of the CA1 hippocampal subfield, corresponding to Schaffer collateral synapses, and the *stratum lucidum* of the CA3 region, representing mossy fiber synapses. High-resolution imaging was performed using an Airyscan detector on a Zeiss LSM 880 confocal microscope equipped with a 63× oil-immersion objective lens (NA 1.4).

Image acquisition involved averaging four-line scans for the Homer1 and Bassoon channels and a single line scan for the DAPI channel, all captured at 16-bit depth. Z-stacks were acquired with a step size of 0.21 μm across a total depth of 8.01 μm.

Laser excitation settings were: 633 nm for Homer1, 561 nm for Bassoon, and 405 nm for DAPI. Photomultiplier tube gain, laser power, and offset settings were optimized during initial calibration and then kept constant across all imaging sessions to ensure consistency.

### Quantification and statistical analysis

Three-dimensional reconstruction of z-stack images for synapse quantification were performed in the Imaris software (Oxford Instruments, Imaris 10.1.1). The “Spots” function was employed to reconstruct Bassoon and Homer1 puncta with a spot size of 0.35 µm, while the “Surfaces” function was used to segment DAPI-stained nuclei. A fixed value threshold was applied uniformly across all sections to ensure consistency in Bassoon and Homer1 spot detection.

To exclude nuclear-associated signals, spots located within DAPI-defined surfaces were removed using the in-built exclusion function. Synaptic colocalization was assessed using the “Colocalize Spots” MATLAB XTension, with a colocalization distance threshold set at 0.35 μm. The total number of colocalized Homer1 and Bassoon puncta, representing putative synapses, was quantified. To compute synapse density, the number of colocalized spots outside nuclear (DAPI) volume was divided by the effective tissue volume (i.e., total region of interest volume minus the DAPI-defined volume). Synaptic density values were averaged across three technical replicates to generate a single value per animal.

All statistical analyses were conducted using GraphPad Prism 11. Independent *t*-tests, Mann–Whitney U tests, and one-way and two-way ANOVAs were employed as specified in the corresponding figure legends. For associations, Pearson’s and Spearman’s correlations were utilized as indicated. Sample sizes are indicated within the bars of each graph and refer to the number of mice used for each experiment. The threshold for statistical significance was *P* < 0.05.

